# Proper names from story recall are associated with beta-amyloid in cognitively unimpaired adults at risk for Alzheimer’s disease

**DOI:** 10.1101/2020.05.22.106195

**Authors:** Kimberly D. Mueller, Rebecca L. Koscik, Lianlian Du, Davide Bruno, Erin M. Jonaitis, Audra Z. Koscik, Bradley T. Christian, Tobey J. Betthauser, Nathaniel A. Chin, Bruce P. Hermann, Sterling C. Johnson

**Affiliations:** Department of Communication Sciences and Disorders, University of Wisconsin-Madison, Madison, WI, USA; Wisconsin Alzheimer’s Institute, University of Wisconsin School of Medicine and Public Health, Madison, WI, USA; Wisconsin Alzheimer’s Disease Research Center, University of Wisconsin School of Medicine and Public Health, Madison, WI, USA; School of Natural Sciences and Psychology, Liverpool John Moores University, Liverpool, UK; Waisman Laboratory for Brain Imaging and Behavior, University of Wisconsin – Madison, Madison, WI, USA; Department of Medical Physics, University of Wisconsin – Madison, Madison, WI, USA; Department of Neurology, University of Wisconsin School of Medicine and Public Health, University of Wisconsin – Madison, Madison, WI, USA; Geriatric Research Education and Clinical Center, William S. Middleton Veterans Hospital, Madison, WI, USA

**Keywords:** Alzheimer’s disease, beta-Amyloid, MCI, Semantic memory, Proper names, Story Recall, Language, Early detection

## Abstract

Due to advances in the early detection of Alzheimer’s disease (AD) biomarkers including beta-amyloid (Aβ), neuropsychological measures that are sensitive to concurrent, subtle changes in cognition are critically needed. Story recall tasks have shown sensitivity to early memory declines in persons with Mild Cognitive Impairment and early stage dementia, as well as in persons with autosomal dominantly inherited AD up to 10 years prior to a dementia diagnosis. However, the evidence is inconclusive regarding relationships between evidence of Aβ and story recall measures. Because story recall tasks require the encoding and delayed retrieval of several lexical-semantic categories, such as proper names, verbs, and numerical expressions, and because lexical categories have been shown to be differentially impaired in persons with MCI, we focused on item-level analyses of lexical-semantic retrieval from a quintessential story recall task, Logical Memory from the Wechsler Memory Scale. Our objective was to investigate whether delayed recall of lexical categories (proper names, verbs and/or numerical expressions), as well as the traditional total score measure, was associated with “preclinical AD,” or cognitively unimpaired adults with positive Aβ deposition on positron emission tomography (PET) neuroimaging using Pittsburgh Compound B. We developed an item-level scoring system, in which we parsed items into lexical categories and examined the immediate and delayed recall of these lexical categories from 217 cognitively unimpaired participants from the Wisconsin Registry for Alzheimer’s Prevention. We performed binary logistic regression models with story recall score as predictor and Aβ status (positive/negative) as the outcome. Using baseline Logical Memory data, proper names from delayed story recall were significantly associated with Aβ status, such that participants who recalled more proper names were less likely to be classified as PiB(+) (odds ratio = .58, p=.01). None of the other story recall variables, including total score, were associated with PiB status. Secondary analyses determined that immediate recall of proper names was not significantly associated with Aβ, suggesting a retrieval deficit rather than that of encoding. The present findings suggest that lexical semantic retrieval measures from existing story recall tasks may be sensitive to Aβ deposition, and may provide added utility to a widely-used, long-standing neuropsychological test for early detection of cognitive decline on the AD continuum.

## 1. Introduction

Alzheimer’s disease (AD) is a neurodegenerative process defined by the presence of beta amyloid plaques (Aβ) and neurofibrillary tau tangles, resulting in neuronal cell death, eventual cognitive decline, and dementia (Braak & Braak, 1991; Jack Jr et al., 2018). Although Alzheimer’s dementia has a heterogenous presentation, the gradual and insidious cognitive decline leading to dementia typically involves documentable changes to both episodic memory, especially the learning of new material, and semantic memory, particularly the timely and efficient word retrieval from a variety of lexical-semantic categories. These patterns of cognitive decline match the evidenced early distribution of pathologic tau, which typically begins in the medial-temporal lobe, particularly the entorhinal cortex which mediates episodic learning and memory, and the parahippocampal gyrus which mediates semantic memory. There is accruing evidence showing that tests of both episodic memory (such as the Rey Auditory Verbal Learning Test (R-AVLT) or Logical Memory story recall from the Wechsler Memory Scale (WMS) (Mormino & Papp, 2018) and tests of semantic memory (such as category verbal fluency or face-name Association tasks) (Rentz et al., 2011) are associated with very early amyloid plaque accumulation, even antecedent to the onset of clinical cognitive impairment.

Because the AD neuropathological processes are suspected to begin decades before clinically significant cognitive decline, interventions are expected to be most effective at the earliest stages of disease, before considerable neurodegeneration has occurred. As a result, sensitive measures of early cognitive change are needed to identify the individuals who will be most likely to benefit from interventions, and as a means of monitoring response to treatment in clinical trials. The National Institute on Aging-Alzheimer’s Association research framework for Alzheimer’s disease indicates that cognitively unimpaired individuals with Aβ *in vivo* biomarker positivity are characterized as in the early stage of Alzheimer’s pathologic change (Jack Jr et al., 2018), and therefore considered to be an ideal group to study for early detection of cognitive decline due to AD and for response to treatment in intervention trials.

Studies evaluating relationships between Aβ deposition using Positron Emission Tomography (PET) imaging and cognition in at-risk, yet otherwise healthy individuals have shown mixed results. While some cross-sectional studies show an association between Aβ and cognition (Donohue et al., 2017), other studies have found no such associations (Jansen et al., 2018).

Longitudinal studies evaluating the change in cognition in cognitively unimpaired adults in relation to Aβ deposition have shown positive relationships, such that Aβ positive individuals tend to show faster prospective declines on cognitive composite scores and on tests of verbal learning and memory than individuals who were Aβ negative (Betthauser et al., 2020; Clark, Racine, et al., 2016; Farrell et al., 2017; Rabin et al., 2018).

Although tests of cognitive function typically focus on episodic memory in AD longitudinal studies, some researchers argue that semantic memory, the long-term storage of conceptual knowledge including meanings and lexical (word) information, may be a more sensitive target for prodromal AD (Venneri, Mitolo, & De Marco, 2016). Because the earliest AD pathology (neurofibrillary tau tangles) and neurodegeneration tend to occur in the entorhinal cortex and parahippocampus (Braak, Braak, & Bohl, 1993), areas known to be neural correlates of semantic processing (Venneri et al., 2016), measurement of semantic processing may be a particularly sensitive target.

Semantic memory deficits have been well documented in AD dementia using a variety of neuropsychological tests, including confrontation naming (Williams, Mack, & Henderson, 1989) category fluency (Garrard, Patterson, Watson, & Hodges, 1998; Rascovsky, Salmon, Hansen, Thal, & Galasko, 2007), and visual-verbal semantic matching (Hodges & Patterson, 1995). Importantly, advances in AD biomarker detection (detection of amyloid plaques and tau neurofibrillary tangles via imaging and cerebrospinal fluid analysis) have allowed researchers to evaluate semantic memory differences between healthy controls and adults in the preclinical phase of AD (i.e., before noticeable functional decline or noticeable symptoms of cognitive impairment). For example, Papp et al. (2016) showed that participants with evidence of elevated Aβ deposition on positron emission tomography (PET) using tracer Pittsburgh Compound B (PiB) showed steeper decline in category fluency than participants without amyloidosis, despite none of the participants in either group met criteria for clinical impairment of cognitive function (Papp et al., 2016).

While category fluency tasks comprise one sensitive way of measuring semantic memory and retrieval, researchers have also found specific lexical categories (i.e., nouns, verbs, proper names) to be differentially affected in typical aging and dementia. For example, proper name retrieval (as compared to regular nouns) is particularly problematic in typical aging (Cohen, 1990) but accentuated in mild dementia due to probable AD (C. Semenza, Borgo, Mondini, Pasini, & Sgaramella, 2000). Lesion studies and functional imaging research have shown that the anterior inferior temporal lobe may be more involved in the retrieval of proper names than other grammatical classes (Gainotti, 2007) and therefore may be sensitive to the early neuropathology associated with AD. Verb disadvantages over nouns have also been documented in AD dementia in tasks such as verb vs. object fluency (Davis et al., 2010) or in action vs. object picture-naming (Druks et al., 2006), but the neural substrates of verb processing are not fully understood. Activation of the dorsolateral prefrontal cortex in verb processing has been documented by several researchers (Cappa, Sandrini, Rossini, Sosta, & Miniussi, 2002; Cotelli et al., 2011; Shapiro, Moo, & Caramazza, 2006), but this region is often spared in the earliest stages of AD. It is possible that association pathways involving semantic memory and retrieval may partially explain these verb deficits (Beber et al., 2019). Perhaps more importantly, the association between verb processing in cognitively unimpaired adults with biomarker evidence of AD pathology is unknown.

In order to explore this question, we capitalized on a widely used memory test, the Logical Memory story recall task from the Wechsler Memory Scale-Revised (Wechsler, 1987). Logical Memory, although an episodic memory test, arguably provides a unique measure of discourse from which variations in semantic categories of spontaneous recall can be culled. In this novel method, we obtained item-level data from participants from the Wisconsin Registry for Alzheimer’s Prevention, a longitudinal cohort study of participants enriched for AD risk who are free of dementia and preclinical conditions (MCI). We separated correctly recalled items into lexical categories including proper names (names of people or places), verbs, and numerical expressions. We had two aims, and hypothesized that recall of either proper names, verbs or numerical expressions would be associated with both 1) progression from cognitively unimpaired to clinical MCI and/or 2) amyloid positivity as evidenced by PiB PET imaging. In sensitivity analyses designed to understand how our novel predictors compared to more traditional measures, we compared the relationship with Aβ status and novel story recall variables to that of a commonly used cognitive composite score, the Preclinical Alzheimer’s Cognitive Composite (PACC) (Donohue et al., 2014).

## 2. Materials and methods

### 2.1 Participants

The study sample was drawn from the Wisconsin Registry for Alzheimer’s Prevention (WRAP) study, an ongoing longitudinal cohort study examining risk factors, lifestyles, and cognition in participants who are late-middle-aged and enriched for parental history of AD. A subset of WRAP participants also participates in AD biomarker studies (e.g., imaging and cerebrospinal fluid studies). The first follow-up visit occurred four years after baseline, and subsequent visits occurred every two years thereafter (see Johnson et al., 2018 for detailed information about the WRAP sample). Logical memory was first added to the test battery in 2007, and this item-level analysis project was added in summer of 2018. Data entry of item-level information was prioritized to focus on those who had provided PET biomarker data and/or had progressed from an unimpaired to impaired status. Once those records were entered, data entry continued for all other charts. At the time of these analyses, participants were selected for these analyses if they were free of dementia at any visit, free of neurological diagnoses (stroke, Parkinson disease, MS, or epilepsy/seizure disorder), had English as their native language, and had their cognitive status reviewed via consensus conference (n=696). A second subset of individuals was selected for the second aim who had completed amyloid PET scans (completed at median visit 3 of WRAP study visit) and met the above-described inclusion criteria (n=217). All activities for this study were approved by the University of Wisconsin – Madison Institutional Review Board and completed in accordance with the Helsinki Declaration.

### 2.2 Experimental variables from Logical Memory story recall

Logical Memory is a story recall subtest of the Weschler Memory Scale – Revised (WAIS-R) (Wechsler, 1987) is a standardized, norm-referenced neuropsychological test that evaluates both learning and episodic memory. Standardized procedures for test administration were followed in accordance with the WMS-R manual. In this task, a first short story (Story A) consisting of a few lines was read aloud to the participant, and the participant was asked to retell the story immediately with the following instructions: “tell the story back to me, using as close to the same words as you can remember; you should tell me all you can, even if you are not sure.” The same procedure was repeated with a second story (Story B). The participant was then asked to recall both stories again after a 25-35 minute delay. Standardized scoring procedures per the WAIS-R manual were followed. Each story contains 25 “idea units”, consisting of a target word, phrase, or idea; as the participant recalled the story, the examiner was instructed to underline correctly expressed idea units, and to notate alternative wordings for scoring later. The scoring criteria allows for some alterations of idea units; for example, the phrase “went off the road” is acceptable for the idea unit “skidded off the road.” Some responses must be verbatim in order to be accepted, including proper names (names of people or places) and numerical expressions (e.g., “two children” or “four days”).

In order to create the lexical category variables, we first set up a database such that each idea unit was a separate variable and was coded as 1 (correctly recalled) or 0 (not recalled). Next, we parsed each idea unit into a lexical category, by running a transcript of each story stimulus through the Computerized Language Analysis Program (CLAN) (MacWhinney, 2014). We obtained the lexical categories of each item using the output of the MOR program in CLAN. Finally, we assigned each idea unit into one of three lexical categories, based on our theory-driven hypotheses: proper names, verbs, and numerical expressions. See **Figure 1** for a schematic of the lexical categories and the idea units they contain (we used a hypothetical story in order to protect copyright and test integrity). For primary analyses, we used data from the first administration of the story recall task as the predictors (median = visit 2, range = visits 1 – 3). Specifically, we summed the semantic categories from stories A and B for each of the immediate and delayed recall conditions resulting in a semantic category total immediate recall, and semantic category total delayed recall. We then converted all variables to standardized z-scores, so that Logical Memory total score and the lexical categories scores were comparable among models.

**Figure 1.**
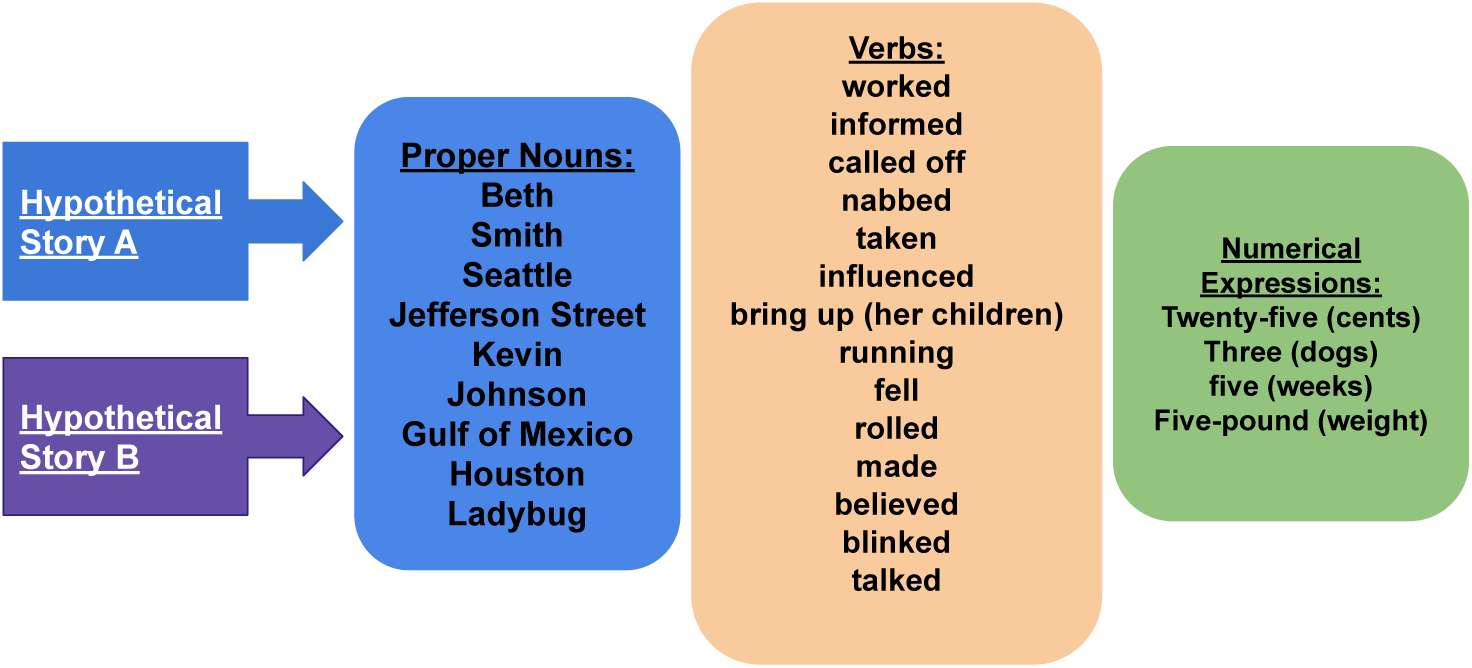
Lexical categories extracted from hypothetical story recall items. Data from this manuscript used Logical Memory from the Wechsler Memory Scale-Revised (Wechsler, 1987) and this schematic is modeled after the Logical Memory subtest in order to protect test integrity/copyright. The above hypothetical schematic represents the same number of items comprising lexical categories as that of the Logical Memory test.

### 2.3 Secondary Predictor: The Preclinical Cognitive Composite Score (PACC)

As first described by Donohue et al. (2014), the PACC is an average of commonly used tests of episodic memory, executive function and global cognition, including total recall from the Free and Cued Selective Reminding Test, delayed recall score from Logical Memory IIa, Digit Symbol Substitution Test, and the MMSE total score. Recently our group compared several versions of the PACC, as well as theoretically derived cognitive composite scores, and found that within the WRAP group a version referred to as “PACC-3” was sensitive to cognitive decline. This version omits the MMSE and includes an average of standardized scores from total recall from Rey AVLT, delayed recall from Logical Memory A & B, and Digit Symbol Substitution (Jonaitis et al., 2019). In secondary analyses, we examined whether the PACC-3 from the first available visit (median visit = 2) would predict morel variability in PiB positivity than the lexical categories variables.

### 2.4 Cognitive Diagnosis

Because the WRAP cohort is relatively young (mean age at baseline = 54; mean age at most recent visit = 67), and the majority of participants are cognitively unimpaired, one priority for the WRAP study has been to develop methods that detect subtle declines in cognition, despite the fact that participants’ scores may still fall within the norm-referenced range of normal. Researchers from our group have therefore developed internal “robust” norms, in which the normative group consists of non-declining WRAP participants over time (Clark, Koscik, et al., 2016; Koscik et al., 2014). Algorithms were developed to “flag” participants whose cognitive performance falls outside the range of the robust norms at any particular visit; flagged participants’ cases are then brought to a consensus review committee consisting of dementia specialists, including neuropsychologists, physicians, nurse practitioners, and physician assistants. The details of this consensus review are described elsewhere (S. Johnson et al., 2018; Koscik et al., 2016); in brief, the committee reviews cognitive data, medical and social histories, mental health status, and self- and informant-reported assessments of mood, cognitive and functional status across all available visits. Clinicians and researchers are blinded to biomarker status prior to making their cognitive status diagnoses. A diagnosis of “clinical MCI” is assigned if the consensus panel agrees that the participant met the criteria for MCI from the National Institute on Aging-Alzheimer’s Association (Albert et al., 2011). The research category “cognitively unimpaired-declining” is assigned to participants who show lower than expected performance (>1.5 standard deviations below internal robust norms), but few or no subjective complaints or functional deficits. This category is similar to clinical stage 2 from the 2018 diagnostic Alzheimer’s disease framework (Jack Jr et al., 2018), but without consideration of biomarker status. Participants who do not meet the above-noted criteria or are not flagged by the algorithm are classified as “cognitively unimpaired-stable.” For the purposes of these analyses, we were interested in those participants who had a diagnosis of “cognitively unimpaired-stable” (CU-S) at the first visit at which story recall was administered, and whether or not they progressed to clinical MCI at their most recent visit.

### 2.5 [^11^C]Pittsburgh compound B PET

For the second research question, participants were selected who underwent a 70-minute dynamic [^11^C]Pittsburgh compound B (PiB) scan on a Siemens EXACT HR+ scanner (and prior to 2015, a T1-weighted magnetic resonance imaging (MRI) scan on a GE 3.0 T MR750 using an 8-channel head coil). Neuroimaging was completed, on average, 1.4 years (SD = 1.4) after the first visit at which story recall was administered (median visit 2). [^11^C]PiB radiosynthesis, acquisition and reconstruction parameters, image processing and quantification have been described previously (S. C. Johnson et al., 2014). Briefly, the PET time series was motion corrected, de-noised, and co-registered to T1-w MRI. Time-activity curves were extracted from the co-registered PET data in subject MRI space from gray matter restricted Anatomic Labeling atlas (AAL, add citation) regions of interest (ROIs) warped to MRI space and used to estimate ROI-level distribution volume ratios (DVRs; Logan Graphical Analysis, cerebellum GM reference region, 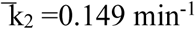; (Logan et al., 1996; Lopresti et al., 2005)). PiB positivity was ascertained by applying a threshold to the mean DVR (global PiB) across eight bilateral ROIs (global DVR ≥ 1.19) (Racine et al., 2016).

### 2.6 Statistical Analyses

All statistical analyses were performed in R (Team, 2019) or SPSS version 25. Linear mixed effects models were run with the R package ‘lme4’ (Bates et al., 2015). Significance level was set at p<.05. To account for multiple comparisons, a false discovery rate approach was applied, which calculates the proportion of false positives among those tests for which the null hypothesis is rejected. This aids in controlling error while preserving power, by adjusting the *p-*value criterion for significance based on the number of tests performed (Curran-Everett, 2000).

#### 2.6.1 Relationship between lexical categories from story recall and progression to clinical MCI

A series of binomial logistic regression analyses were performed to model the relationship between progression to clinical MCI at the most recent visit (progression = 1; no progression = 0) and each of four story recall scores from the first available visit, considered in separate models: the number of words recalled in each lexical category, and the total number recalled. Covariates included age, sex, and Wide-Range Achievement Test – 3 (WRAT-3)) (Wilkinson, 1993) reading subtest were included in each model. We use the WRAT-3 reading test as a proxy for educational attainment as described elsewhere (Manly, Jacobs, Touradji, Small, & Stern, 2002). Within each set of models, we used the Akaike information criterion (AIC) to assess whether any of the Logical Memory lexical sub-scores provided superior fit to that obtained using the traditional total score. The AIC is an estimator of the relative quality of statistical models, with the preferred model containing the minimum AIC (Bozdogan, 1987). In order to determine if the model fits were comparable, we followed guidance from Burnham & Anderson (2002), such that models were deemed comparable when the difference in AIC relative to the smaller AIC model was < 2 (Burnham & Anderson, 2002).

#### 2.6.2 Relationship between lexical categories and Aβ positivity

Similar binomial logistic regression analyses were performed to model the relationship between amyloid status at the most recent PET scan (PiB(+) = 1; PiB(-) = 0) and the lexical category scores and total scores. Covariates included age, sex, WRAT-3 and *APOE-ε4* status (*APOE-ε4* positive = at least one allele). Primary analyses focused on delayed recall scores in order to capture retrieval from semantic memory vs. encoding; immediate recall scores were secondary. Again, AICs were compared between models to identify the best fitting model(s). In sensitivity analyses, we repeated the analysis substituting the Logical Memory variables from the most recent visit for the baseline Logical Memory variables. In additional exploratory analyses, we examined 1) whether models including multiple lexical variables or a lexical variable and remaining total score predicted more variability than just single lexical variables and covariates; and 2) whether there was a relationship between longitudinal story recall variables and Aβ at most recent PET scan.

#### 2.6.3 Bootstrapping to determine the significance of association and the stability of findings

We applied the following bootstrapping technique to approximate a distribution and test the stability of regression coefficient estimates for each of the eight logistic regression models in 2.5.2. First, we resampled the first available visit data with replacement to obtain 1000 samples the same size as our analysis sample. We then ran the batch of logistic regression models and stored the coefficients from each. We repeated these two steps 200 times. We used the resulting distributions of parameter estimates to obtain 95% confidence intervals.

## 3. Results

### 3.1 The relationship between lexical categories and progression to clinical MCI

Participant demographics and clinical characteristics for the first study, the relationship between lexical categories and progression to clinical MCI, are presented in Table 1. 696 participants had a mean age at visit 2 of 58 (sd = 6) and mean age at most recent visit was 65 (sd=6.8). 70% were female, 90% were non-Hispanic white, and 76% had a parental family history of Alzheimer’s disease dementia. Eighteen of the 696 participants (2.6%) progressed from cognitively unimpaired-stable at the first story recall visit (median = visit 2) to clinical MCI at the most recent visit (median = visit 5). Those who progressed to MCI did not differ from those who were unimpaired at their last visit by race, parental history of AD, WRAT-3 Reading, years of education, APOE-ε4 status, or Mini-mental state examination scores from visit 2. The participants who progressed to MCI were significantly older than the CU-S participants (64 vs. 58 at first available story recall, p <.001)), and tended to have a higher percentage of males (56% of participants with MCI were male vs. 30% of CU-S participants, p = .04). The MCI participants performed significantly worse on Rey-AVLT, Logical Memory total score, and all Logical Memory experimental lexical categories at visit 2, adjusting for age, sex and WRAT-3 reading (when all participants in this sample were deemed cognitively unimpaired – stable).

**Table 1.**
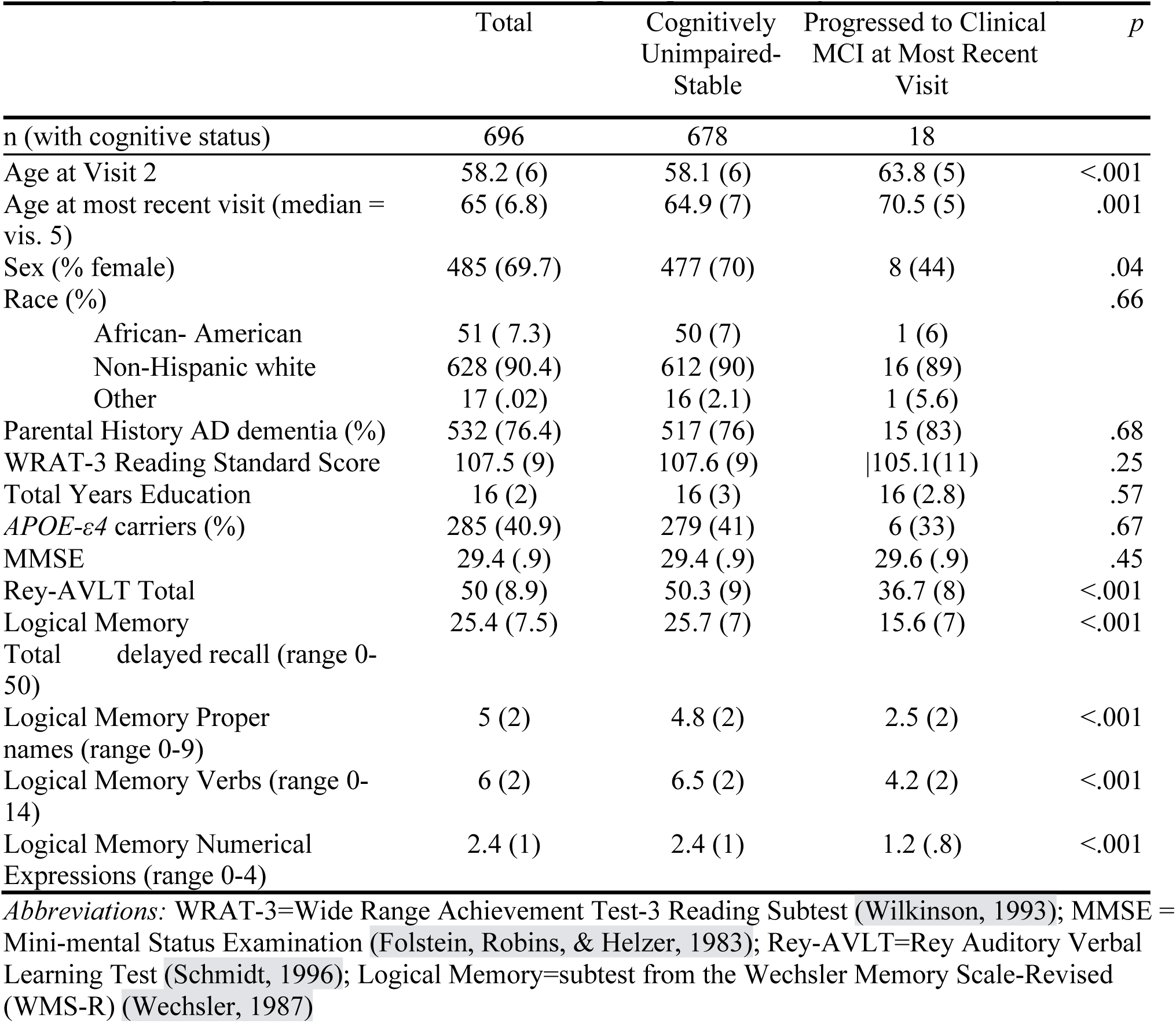
Demographics and clinical characteristics of participants with cognitive status and story recall

Results of the logistic regression analyses using story recall delayed total score and lexical categories as predictors for conversion to clinical MCI are presented in Table 2. All variables from story recall (proper names, verbs, numerical expressions and total score) were significant predictors of conversion to clinical MCI at the most recent visit. When comparing the AICs among all four models, the model using total score as predictor was the better fitting model (AIC=38.36); however, the four models’ AICs did not differ by more than 2, indicating similar fits overall (Burnham & Anderson, 2002).

**Table 2.**
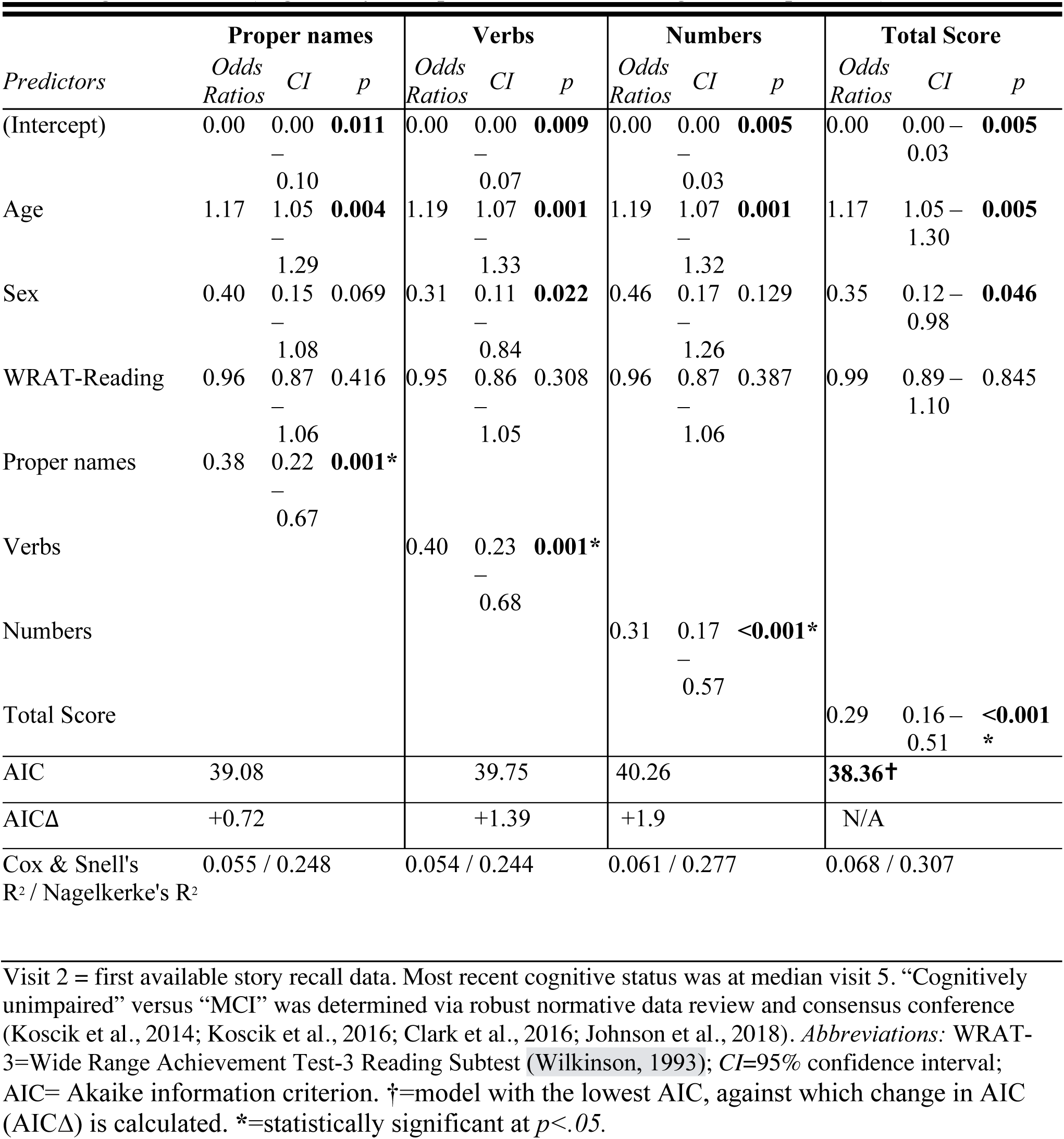
Binomial logistic regression results of story recall variables at visit 2 predicting most recent cognitive status (cognitively unimpaired versus Mild Cognitive Impairment)

### 3.2 The relationship between lexical categories and beta-amyloid status

Participant demographics and clinical characteristics of participants with PET-PiB imaging data are presented in Table 3. At the time of these analyses, a total of 217 participants had both PiB data and coded story recall variables. 172 participants were classified as PiB(-) and 45 were PiB(+) at the most recent imaging visit. The PiB(+) group was significantly older (mean age = 61, sd=5) than the PiB(-) group (mean age = 58, sd=6) (p=.005), and had a higher percentage of APOE-e4 carriers (72% in the PiB(+) group vs. 31% in the PiB(-) group). The two groups did not differ in any other variables, including education, WRAT-3 reading scores, Mini-Mental State Examination scores, or Rey-AVLT total scores.

**Table 3.** Demographics and clinical characteristics of subset of WRAP participants with PiB-PET imaging

|  | PiB(-) | PiB(+) | p |
| --- | --- | --- | --- |
| n | 172 | 45 |  |
| Age at visit 2 (mean (sd)) | 58.11 (6.22) | 60.96 (4.86) | **0.005** |
| Age at PET Scan | 65.3 (6.8) | 69 (5.3) | **<.001** |
| Age at PET – Age at visit 2 | 7.2 (3) | 8.2 (3) | **0.04** |
| Female (%) | 121 (70.3) | 30 (66.7) | 0.767 |
| Progressed to clinical MCI = 1 (%) | 3 (1.7) | 2 (4.4) | 0.605 |
| race (%) |  |  | 0.813 |
| Native American | 3 (1.7) | 0 (0.0) |  |
| Asian | 1 (0.6) | 0 (0.0) |  |
| African-American | 5 (2.9) | 2 (4.4) |  |
| Non-Hispanic White | 162 (94.2) | 43 (95.6) |  |
| Other | 1 (0.6) | 0 (0.0) |  |
| Family History positive (%) | 123 (71.5) | 36 (80.0) | 0.339 |
| WRAT-3 Reading (mean (sd)) | 108.56 (8.50) | 109.25 (7.63) | 0.626 |
| Total Years of Education (mean (sd)) | 16.38 (2.77) | 17.22 (2.91) | 0.073 |
| **APOE carrier (%)** | 56 (32.6) | 32 (71.1) | **<0.001** |
| Baseline MMSE (mean (sd)) † | 29.37 (0.96) | 29.40 (0.84) | 0.858 |
| Baseline R-AVLT Total Score (mean (sd)) † | 55.6 (7.8) | 54.6 (7.9) | 0.416 |
| Baseline Logical Memory delayed total (mean (sd)) † | 26.34 (7.13) | 26.91 (6.66) | 0.630 |
| Proper Name delayed total | 5.3 (2.0) | 4.4 (1.9) | **0.010** |
| Verb delayed total | 6.6 (2.2) | 6.2 (1.3) | 0.214 |
| Numerical Expressions delayed total | 2.4 (1.1) | 2.6 (1.1) | 0.520 |

†"baseline"=collected at the first visit at which Logical Memory was administered, median=visit 2. Items in **bold** are statistically significant at $p<.05$ using t-tests or chi-square tests (unadjusted means).
Abbreviations: WRAT-3=Wide Range Achievement Test-3 Reading Subtest (Wilkinson, 1993); MMSE = Mini-mental Status Examination (Folstein, Robins, & Helzer, 1983); Rey-AVLT=Rey Auditory Verbal Learning Test (Schmidt, 1996); Logical Memory=subtest from the Wechsler Memory Scale-Revised (WMS-R) (Wechsler, 1987)

Results from the logistic regression models are presented in Table 4. Using baseline Logical Memory data, proper names from delayed story recall were significantly associated with PiB status, such that participants who recalled more proper names from the stories were less likely to be classified as PiB(+) (odds ratio = .58, p=.01). None of the other story recall variables, including total score, were associated with PiB status, and the proper names model was the best fitting model when the AICs were compared; specifically, the proper names model’s AIC was 5.63 units lower than the next best fitting model (verbs), which surpasses the criteria of > 2 units indicating better fit (Burnham & Anderson, 2002).

**Figure 2** provides a supporting visual illustration of %PiB(+) by low, medium, and high z-scores for delayed recall of proper nouns (left) and delayed total recall (right).

**Figure 2.**
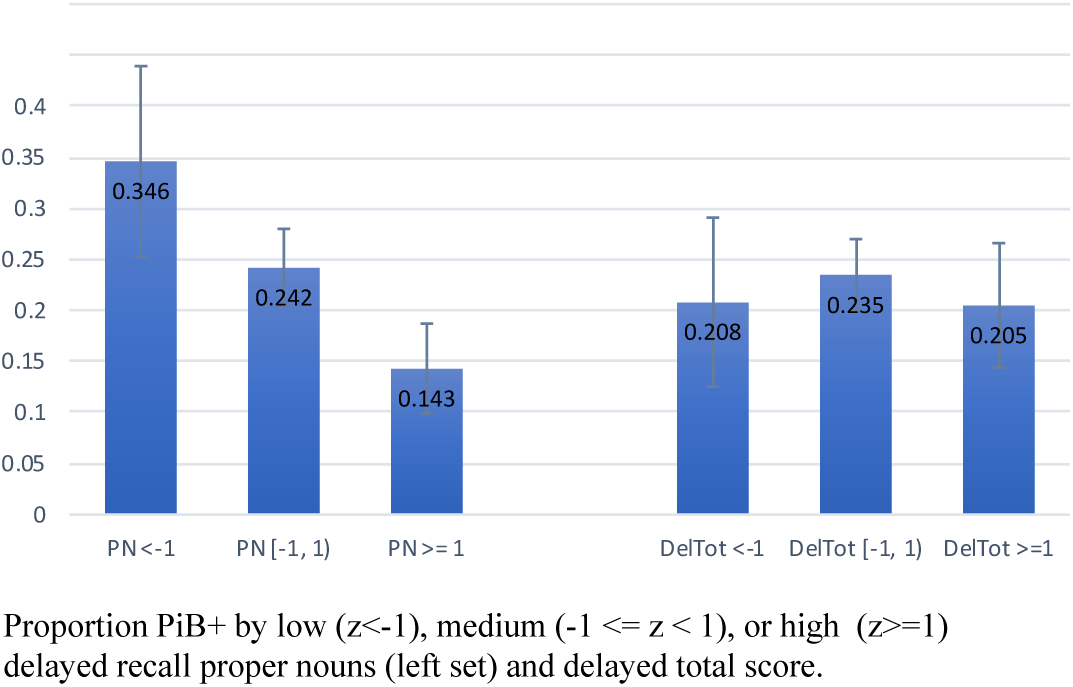
Percent amyloid positive by low, medium and high z-scores for delayed recall of proper names (left) and delayed recall total (right). Proportion PiB+ by low (z<-1), medium (−1 <= z < 1), or high (z>=1) delayed recall proper nouns (left set) and delayed total score.

**Table 4.** Binomial logistic regression results of story recall variables at visit 2 predicting most recent beta-amyloid status

| *Predictors* | **Proper names** |  |  | **Verbs** |  |  | **Numbers** |  |  | **Total Score** |  |  |
| --- | --- | --- | --- | --- | --- | --- | --- | --- | --- | --- | --- | --- |
|  | *Odds Ratios* | *CI* | *p* | *Odds Ratios* | *CI* | *p* | *Odds Ratios* | *CI* | *p* | *Odds Ratios* | *CI* | *p* |
| (Intercept) | 0.00 | .00 – .00 | **<0.001** | 0.00 | 0.00 – 0.00 | **<0.001** | 0.00 | 0.00 – 0.00 | **<0.001** | 0.00 | 0.00 – 0.00 | **<0.001** |
| Age at PiB | 1.13 | 1.05 – 1.22 | **0.001** | 1.15 | 1.07 – 1.23 | **<0.001** | 1.14 | 1.06 – 1.23 | **<0.001** | 1.14 | 1.06 – 1.23 | **<0.001** |
| PiB Age-Story Recall Age | 1.08 | 0.94 – 1.25 | 0.289 | 1.02 | 0.89 – 1.18 | 0.754 | 1.03 | 0.90 – 1.19 | 0.653 | 1.03 | 0.90 – 1.19 | 0.646 |
| Sex | 1.26 | 0.55 – 2.89 | 0.590 | 0.94 | 0.42 – 2.10 | 0.875 | 0.95 | 0.42 – 2.14 | 0.895 | 0.97 | 0.43 – 2.18 | 0.945 |
| *APOE-ε4* | 9.12 | 3.88 – 21.43 | **<0.001** | 8.73 | 3.83 – 19.91 | **<0.001** | 8.08 | 3.58 – 18.22 | **<0.001** | 7.91 | 3.53 – 17.73 | **<0.001** |
| Proper names | 0.58 | 0.38 – 0.88 | **0.004\*** |  |  |  |  |  |  |  |  |  |
| Verbs |  |  |  | 1.49 | 0.97 – 2.29 | 0.070 |  |  |  |  |  |  |
| Numerical Expressions |  |  |  |  |  |  | 1.20 | 0.80 – 1.80 | 0.371 |  |  |  |
| Total Score |  |  |  |  |  |  |  |  |  | 1.15 | 0.75 – 1.76 | 0.529 |
| Cox & Snell's $R^2$ / Nagelkerke's $R^2$ | 0.206 / 0.322 | | | 0.194 / 0.303 | | | 0.184 / 0.288 | | | 0.182 / 0.285 | | |
| AIC | **175.26†** |  |  | 180.89 |  |  | 183.44 |  |  | 183.8 |  |  |
| AICΔ | N/A |  |  | +5.63 |  |  | +8.18 |  |  | 8.54 |  |  |

*Abbreviations:* WRAT-3=Wide Range Achievement Test-3 Reading Subtest (Wilkinson, 1993); *CI*=95% confidence interval; AIC= Akaike information criterion. †=model with the lowest AIC, against which change in AIC (AICΔ) is calculated; AICΔ=Change in AIC against the model with the lowest AIC, in this case the Proper Names model. \*=statistically significant at False Discovery Rate critical value of $p<.0125$ .

In order to test the robustness of the observed relationship between baseline proper names and most recent PiB status, we ran sensitivity analyses using the logistic regression with most recent proper names recall as predictor (median = visit 4, range = 2-6). We found the same pattern, with proper names significantly predicting PiB status (odds ratio = .64, confidence interval = .45-.92, *p*=.02), while total score was not significantly associated with PiB status. These results were also confirmed when we performed bootstrapping analyses (Table 5). Furthermore, we ran the models with immediate recall of proper names and there was no significant relationship with PiB status.

**Table 5.** Follow up: Binomial logistic regression bootstrapped results of story recall variables at visit 2 predicting most recent beta-amyloid status

|  | Proper Noun |  | Verbs |  | Number |  | Total Score |  |
| --- | --- | --- | --- | --- | --- | --- | --- | --- |
| *Predictors* | *Odds Ratios* | *CI* | *Odds Ratios* | *CI* | *Odds Ratios* | *CI* | *Odds Ratios* | *CI* |
| Age at PiB | 1.19 | 1.16–1.23 | 1.18 | 1.15–1.22 | 1.18 | 1.15–1.22 | 1.18 | 1.15–1.22 |
| WRAT-3 Reading | 1.05 | 0.98–1.11 | 0.99 | 0.93–1.06 | 1.01 | 0.94–1.07 | 1.01 | 0.94–1.07 |
| Sex(female) | 1.69 | 1.17–2.50 | 1.20 | 0.78–1.77 | 1.21 | 0.83–1.78 | 1.26 | 0.87–1.82 |
| *APOE-ε4* | 11.63 | 7.86–19.54 | 9.70 | 6.46–15.56 | 9.25 | 6.12–14.73 | 9.09 | 5.98–14.51 |
| Total Years of Education | 1.17 | 1.04–1.33 | 1.15 | 1.03–1.32 | 1.15 | 1.03–1.31 | 1.15 | 1.02–1.32 |
| Proper names | **0.52** | 0.43–0.64 |  |  |  |  |  |  |
| Verbs |  |  | 1.47 | 1.18–1.99 |  |  |  |  |
| Numerical Expressions |  |  |  |  | 1.16 | 0.96–1.42 |  |  |
| Total Score |  |  |  |  |  |  | 1.05 | 0.85–1.42 |

*Abbreviations:* WRAT-3=Wide Range Achievement Test-3 Reading Subtest (Wilkinson, 1993); *CI*=95% confidence interval. Items in **bold** indicate statistical significance at $p<.05$ .

In exploratory analyses using the baseline delayed Logical Memory data, we examined whether including multiple lexical variables simultaneously explained additional variance in PiB status. After adjusting for proper names, higher verbs and total score were associated with increased risk of amyloid (odds ratio 2.04, p=.0004 and 2.40, p=.008 respectively). Follow-up analyses to understand these patterns indicated that low proper names (z<0) and higher verbs or higher total score (z>0) had higher risk of amyloid positivity than those who were higher on both proper names and verbs or proper names and total (Table 6). We created a 4-level variable via combinations of low vs. high proper nouns and verbs based on median split of each, with the reference group being those with higher counts of proper names and higher counts of verbs. These sensitivity analyses sought to examine whether the results observed using continuous proper name and verb data persisted when simplified to categorical levels.

**Table 6.** Follow-up multinomial logistic regression results comparing risk of amyloid positivity in proper name X verb recall groups.

| Proper Name/Verb Group |  |  |  |  |
| --- | --- | --- | --- | --- |
| *Response\** | *Predictors* | *Odds Ratios* | *CI* | *p* |
| 2 | Age at PiB | 0.97 | 0.91 – 1.04 | 0.433 |
|  | WRAT-3 Reading | 0.89 | 0.80 – 0.99 | **0.039** |
|  | Sex(female) | 0.28 | 0.12 – 0.67 | **0.004** |
|  | *APOE-ε4* | 0.70 | 0.27 – 1.80 | 0.465 |
|  | PiB Status (positive) | 3.81 | 1.32 – 11.02 | **0.013** |
| 3 | Age at PiB | 0.98 | 0.92 – 1.04 | 0.504 |
|  | WRAT-3 Reading | 0.88 | 0.80 – 0.97 | **0.010** |
|  | Sex(female) | 0.68 | 0.28 – 1.64 | 0.390 |
|  | *APOE-ε4* | 2.18 | 0.97 – 4.90 | 0.058 |
|  | PiB Status (positive) | 0.57 | 0.18 – 1.78 | 0.330 |
| 4 | Age at PiB | 1.01 | 0.95 – 1.07 | 0.853 |
|  | WRAT-3 Reading | 0.83 | 0.75 – 0.91 | **<0.001** |
|  | Sex(female) | 0.35 | 0.15 – 0.80 | **0.013** |
|  | *APOE-ε4* | 1.31 | 0.56 – 3.03 | 0.536 |
|  | PiB Status (positive) | 1.11 | 0.38 – 3.23 | 0.841 |
|  | Observations | 216 |  |  |
|  | R<sup>2</sup> Nagelkerke | 0.219 |  |  |

4-level variable created via combinations of low vs. high proper names and verbs based on median split of each. Reference group = high nouns and high verbs. \*Response 1: Noun positive & verb positive (n=81); Response 2: Noun negative, verb positive (n=38); Response 3: Noun positive, verb negative (n=49); Response 4: Noun negative, verb negative (n=49)

### 3.3 Sensitivity analysis: comparing the relationship with beta-amyloid status and proper names to the Preclinical Alzheimer’s Cognitive Composite (PACC)

In additional sensitivity analyses, we also examined whether the proper names model would explain more variability in PiB(+) status than the WRAP version (Jonaitis et al., 2019) of a commonly used cognitive composite score, the Preclinical Alzheimer’s Cognitive Composite (PACC) (Donohue et al., 2014). The PACC-3 composite score from visit 2 was not associated with the most recent PiB status. The AIC for the Proper Name model was 175.26, while the AIC for the PACC-3 model was 189.73, indicating the proper name model was a better fitting model.

### 3.5 Secondary analysis: evaluating longitudinal change in proper names expression and beta-amyloid status

Because of the longitudinal relationships between PiB status and cognitive decline evidenced in the literature, we used linear mixed effects models with either total score or proper names expression as the outcome, and the interaction between PiB status and time as the predictor of interest. Results for these models are presented in Table 7. The interaction between PiB status and age was not significantly associated with proper names recall over time, but this interaction term was significantly associated with total score (*p =* .*02)*. As noted in Table 3, participants who were PiB(+) were already lower on proper names at the baseline story recall visit than those who were PiB(-).

**Table 7.** Results from linear mixed effects model with amyloid status predicting proper names vs. total score over time.

| *Predictors* | **Proper names** |  |  | **Total Score** |  |  |
| --- | --- | --- | --- | --- | --- | --- |
|  | *Estimates* | *CI* | *p* | *Estimates* | *CI* | *p* |
| Age | -0.07 | -0.10 – -0.04 | **<0.001** | -0.20 | -0.33 – -0.08 | **0.001** |
| Sex (female) | 0.56 | 0.13 – 0.99 | **0.011** | 1.94 | 0.34 – 3.53 | **0.017** |
| Practice | 0.06 | -0.05 – 0.17 | 0.263 | 0.67 | 0.30 – 1.03 | **<0.001** |
| WRAT-3 | 0.12 | 0.08 – 0.17 | **<0.001** | 0.55 | 0.39 – 0.72 | **<0.001** |
| PiB Status (positive) | 2.43 | -1.28 – 6.15 | 0.199 | 12.89 | 1.68 – 24.10 | **0.024** |
| PiB * Age | -0.04 | -0.10 – 0.01 | 0.138 | -0.21 | -0.38 – -0.03 | **0.020** |
| **Random Effects** |  |  |  |  |  |  |
| $\sigma^2$ | 2.08 | | | 15.82 | | |
| $\tau_{00}$ | 1.81<sub>WRAPNo</sub> | | | 28.10<sub>WRAPNo</sub> | | |
| ICC | 0.46 |  |  | 0.64 |  |  |
| N | 232<sub>WRAPNo</sub> |  |  | 232<sub>WRAPNo</sub> |  |  |
| Observations | 925 |  |  | 925 |  |  |
| Marginal R<sup>2</sup> /<br>Conditional R<sup>2</sup> | 0.153 / 0.547 |  |  | 0.174 / 0.702 |  |  |

Linear mixed effects model with centered age as the time variable, and the interaction between amyloid status (PiB) and centered age as the predictor of interest.

The predicted values of the simple age slopes for participants who were PiB (+) versus those who were PiB(-) for each model are depicted in **Figure 3**.

**Figure 3.**
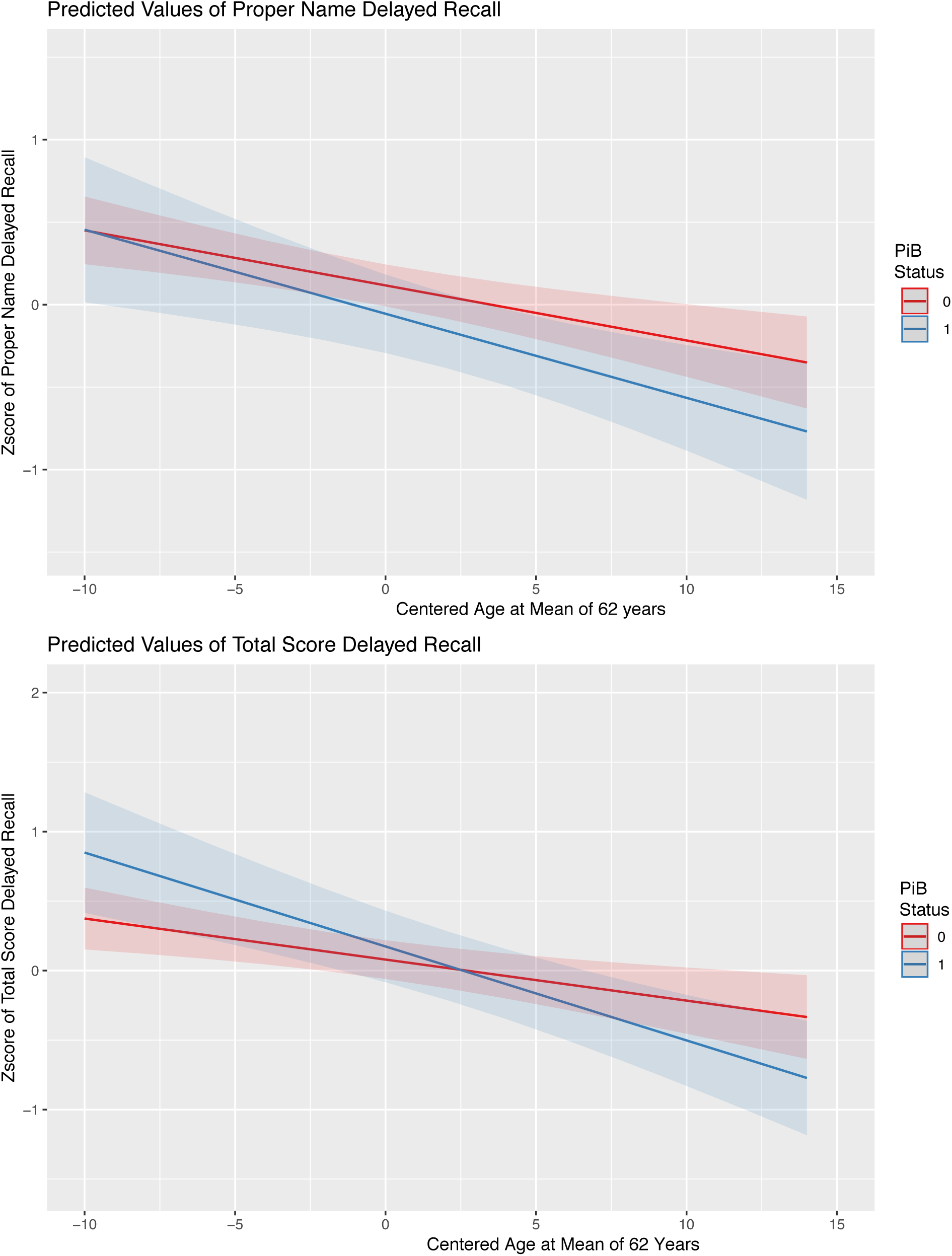
Simple slopes of linear mixed effects models of proper names and total recall as outcomes, and amyloid status (PiB) as the predictor of interest. PiB Status: 0 = amyloid negative, 1=amyloid positive. Age is centered at a mean of 63.4 for ease of interpretation.

## 4. Discussion

At the time of this study, the ability to detect AD neuropathological processes *in vivo*, particularly *both* Aβ plaque accumulation and tau neurofibrillary tangles, is a relatively new advancement in the field of AD research. As a result, the temporal patterns of cognitive decline, or the presence of cognitive differences in those individuals with accumulating AD biomarkers, is largely unknown. In this study, we show a novel finding that cognitively unimpaired participants who were amyloid positive using the Pittsburgh Compound-B (PiB) amyloid PET tracer were less likely to recall proper names from a story recall task at their baseline visit—an average of 7 years earlier—than those without increased PiB accumulation. We did not find such cross-sectional associations with PiB and story recall total score, or with the Preclinical Alzheimer’s Cognitive Composite (PACC). We replicated this finding in two follow-up analyses: one using participants’ most recent visit (an average of 2 years before PiB scan) and the other using bootstrapped analyses from multiple visits. Moreover, this relationship between proper names and PiB positivity was only significant for delayed recall, not immediate recall, thus indicating that it is *retrieval* of proper names, not encoding, that is associated with PiB status.

Proper name retrieval has been shown to be particularly difficult not only in the face of cognitive impairment, but also in typical aging. Several theories of proper name retrieval converge on the basic idea that proper names do not have the network of semantically related attributes to aid in retrieval as regular nouns do (Cohen & Burke, 1993; C. Semenza et al., 2000; Carlo Semenza, Mondini, Borgo, Pasini, & Sgaramella, 2003). It is this limited bank of cognitive resources that retrieval of proper names has to draw upon that may have particular sensitivity to either focal or diffuse brain pathology. Furthermore, the temporal lobe seems more involved in the retrieval of proper names than other lexical categories (Gorno-Tempini et al., 1998), also corresponding to the typical location of early pathologic tau accumulation (Braak & Braak, 1991). Our findings here are supported by other studies that include formal tests of proper name retrieval, such as face-name association tasks or recall of recent and remote famous names. For example, Orlovsky et al. (2017) examined the remote and recent famous face-name recall in a group of unimpaired older adults (mean age = 78) who were either Aβ+ or Aβ-. The Aβ+ group recalled fewer recent proper names than the Aβ-group, even when provided with phonological (letter) cues (Orlovsky et al., 2018). Rentz et al. (2011) found that a demanding face-name association task was more sensitive to Aβ pathology than the 6-Trial Selective Reminding Test in a group of cognitively unimpaired older adults with a mean age of 71 (Rentz et al., 2011).

Our study is different from these previous findings about proper names and early amyloid accumulation in several ways. First, the WRAP subset of participants with amyloid imaging and longitudinal neuropsychological testing is younger than most of the groups studied previously, with a mean age of 66 at PiB scan, and a mean age of 58 at the baseline story recall visit. Second, our proper name findings stem from a novel subcomponent of a commonly used test of episodic memory: story recall. To our knowledge, this is the first study to examine lexical categories from story recall and to show that there may be added sensitivity to Aβ accumulation above and beyond total score. This fact carries special importance in the field today, as there are multiple, ongoing and worldwide longitudinal studies of at-risk cohorts administering story recall, including studies in the Alzheimer’s Disease Neuroimaging Initiative (ADNI), studies in the National Alzheimer’s Coordinating Center (NACC), and the European Alzheimer’s Disease Consortium. A search using the term “Logical Memory” in the Global Alzheimer’s Association Interactive Network (GAAIN) yields well over 36,000 participants who have completed this test for an Alzheimer’s disease research study. Because adding new tests to existing longitudinal studies is a concern due to time constraints, understandable reluctance to add to participant burden, and the need for increased staff time and resources, the prospect of using existing data in new ways is invaluable.

Although cross-sectional associations with proper name recall and Aβ status were present across multiple time points, significantly worse longitudinal *change* in proper names recall over time was not associated with Aβ status. There are several possibilities that may explain this disparity. First, delayed recall of proper names at baseline differed between PiB+ and PiB-while delayed total recall did not. Furthermore, the range of proper names from stories A and B from Logical memory is small, from 0-9 (versus total score which ranges from 0-50). Thus, it is possible that participants showing subtle cognitive impairment associated with Aβ were already low at baseline logical memory and had less room to decline. That is, participants may recall very few proper names very early on, and perhaps an even earlier starting point – before amyloid accumulation reaches positivity – is required to see such change in proper name retrieval. Recent work by our group and others has shown that by the time a person has reached a PET amyloid positivity threshold, brain amyloid has already been accumulating for many years (Koscik, Betthauser, et al., 2020). Future analyses will examine whether duration of PiB at cognitive baseline predicts story recall components or decline in story recall variables.

Secondary analyses from our study showed that when verbs and total score were included in the proper name model, participants who recalled fewer proper names but higher numbers of verbs or total score were more likely to be classified as PiB positive than those who recalled higher proper names and higher verbs or total score. This finding may represent an overcompensation, such that subtle word retrieval problems may result in circumlocution and result in more talking overall. Because the Logical Memory scoring manual allows alternate wording for certain items in verbs and other lexical categories that make up the total score (i.e., using our hypothetical example, the word “cancelled” would be awarded a point for the words “called off, “or “spoke” would be an allowed alternate response for “talked”), such circumlocution and non-verbatim responses might result in a greater total score. Conversely, the proper names in each of the stories require verbatim or near-verbatim responses (e.g., “Sue” would be awarded a point for “Suzy”). Future analyses for understanding these dissociations (low proper names but high total scores) may include examining proportions of total score, as well as performing discourse analyses on digitally recorded story recall in order to quantify behaviors such as circumlocutions, order of recall, and errors and error monitoring.

This study also examined whether proper name recall explained additional variability above and beyond total score in terms of which participants progressed to Clinical MCI from cognitively unimpaired. Although all story recall variables significantly contributed to this prediction, proper names did not explain additional variability. A limitation of the clinical progression analysis was its circularity: story recall total score is a factor used in determining cognitive impairment in the WRAP consensus process. In future analyses, we can use actuarial definitions of impairment (Bondi et al., 2014; Jak et al., 2016) based on tests not including Logical Memory and our internally derived norms (Clark, Koscik, et al., 2016; Koscik et al., 2019; Koscik et al., 2014) to compare how the lexical categories predict progression relative to the traditionally-used total scores.

### 4.1 Strengths and Limitations

This study presents with strengths and limitations which should be acknowledged. First, as noted previously, a significant strength is the use of existing data in a novel way, especially due to the fact that there is a wealth of this data worldwide that can be used to attempt to replicate our findings. Second, the nature of the story recall task itself is a strength; that is, the act of retelling a story that includes proper names, events and other details is a closely related snapshot of an everyday activity in which people engage. This ecological validity can help to uncover the real-world problems that people with subtle cognitive impairments face in communicating, and can inform both pharmacological and nonpharmacological intervention studies. Last but not least, the inclusion of individuals with PET imaging data confirming the presence of amyloid pathology allowed us to determine the relationship between lexical category retrieval and pathology, as opposed to clinical or sub-clinical findings, which are often variable from visit to visit, and may have myriad underlying causes. This study allows the interpretation that lexical retrieval may be related to preclinical AD, although replication in other cohorts and with additional longitudinal study visits and biomarker data is necessary.

A limitation of this study is the sample itself. WRAP is a self-selected, family history cohort made up of participants largely residing in the Upper Midwest. Participants generally are highly educated, and the sample is predominantly non-Hispanic White, thus not representative of the general population. This underscores the need for replication in other cohorts to confirm these associations. As noted above, an additional limitation exists in the ability to use these measures to understand progression to clinical status due to the circular nature of using subcomponents of a test widely used to determine cognitive decline.

### 4.2 Conclusion

Our data suggest an early association between delayed recall of proper names from a story recall task and beta-amyloid accumulation. The wealth of story recall data that exists in longitudinal at-risk cohort studies will allow for possible replication of these findings, without adding additional participant burden. Future directions for this work include examining these associations with evidence of other AD biomarkers from imaging and cerebrospinal fluid, including markers of phosphorylated tau and neurodegeneration, and in other cohorts. Examining correlations between regional atrophy and proper name retrieval will also contribute understanding of these results.

